# OPUS-DSD: Deep Structural Disentanglement for cryo-EM Single Particle Analysis

**DOI:** 10.1101/2022.11.22.517601

**Authors:** Zhenwei Luo, Fengyun Ni, Qinghua Wang, Jianpeng Ma

## Abstract

Many Cryo-EM datasets contain structural heterogeneity due to functional or nonfunctional dynamics that conventional reconstruction methods may fail to resolve. Here we propose a new method, OPUS-DSD (Deep Structural Disentanglement), which can reliably reconstruct the structural landscape of cryo-EM data by directly translating the 2D cryo-EM images into 3D structures. The method adopts a convolutional neural network and is regularized by a latent space prior that encourages the encoding of structural information. The performance of OPUS-DSD was systematically compared to a previously reported method, cryoDRGN, on synthetic and real cryo-EM data. It consistently outperformed existing methods, resolved large or small structural heterogeneity, and improved the final reconstructions of tested systems even on highly noisy cryo-EM data. The results have shown that OPUS-DSD should be particularly suitable for cases in which the high structural flexibilities cannot easily be represented by rigid-body movements. Therefore, OPUS-DSD represents a valuable tool that can not only recover functionally-important structural dynamics missed in a traditional cryo-EM refinement, but also improve the final reconstruction by increasing homogeneity in a dataset. OPUS-DSD is available at https://github.com/alncat/opusDSD.

## Introduction

Macromolecules carry out their functions via specialized motions. Structural biology aims to not only determine the three-dimensional (3D) structures of macromolecules to high resolutions, but also understand their functions by recovering structural dynamics. Cryo-electron microscopy (Cryo-EM) single particle analysis is a powerful structure determination method for obtaining high-resolution 3D structures (1–3). However, the conventional cryo-EM reconstruction in general only produces a single static 3D model without dynamic information. The large numbers of snapshots captured by cryo-EM preserve a huge amount of heterogeneity originated from either conformational or compositional variations or both in 3D structures. Therefore, the structural heterogeneity for macromolecules can be properly revealed if new powerful analysis methods are available (1). Resolving structural heterogeneity would also help improve the resolution of final 3D model in cryo-EM single particle analysis.

The conventional tool for resolving heterogeneity in cryo-EM datasets is 3D classification (4, 5), which models different conformations as individual 3D volumes. The 3D classification scales poorly with respect to (w.r.t) the number of conformations. Additionally, 3D classification can only detect significant structural changes as it assumes the dataset has a few major conformations. Therefore, 3D classification fell short of resolving structural dynamics composed of a large number of transitional conformations. Other approaches like multibody refinement in RELION (6) and 3D variability analysis (3DVA) in cryoSPARC (7) model continuous conformations changes using linear combinations of reaction coordinates. The expression power of these methods is limited to the linear dynamics of a consensus model. Moreover, they are unable to model the discontinuous compositional changes in cryo-EM datasets.

CryoDRGN pioneered a new direction of resolving structural heterogeneity using neural network as a representation for 3D structures (8). Their approach provides a unified framework for resolving structural heterogeneity ranging from conformational to compositional changes. Resolving structural heterogeneity has been formulated as learning a neural network which translates 2D images to its 3D structures directly in cryoDRGN (8). Though cryoDRGN has demonstrated its success in reconstructing an ensemble of 3D structures from consensus refinement result, there are still many challenges that are inherent to its formulation and limit its structural resolving power. In this paper, we present a new method, OPUS-DSD (deep structural disentanglement), for 3D structural heterogeneity analysis.

The landscape of 3D structures is assumed to admit a low dimensional latent representation. Translating 2D image to 3D structure is equivalent to mapping the 2D image to the encoding of its underlying 3D structure in latent space using neural networks, which can also perform the reverse task, the decoding of latent code to 3D structure. The continuous transformation between 2D image and 3D structure can then be learned end-to-end. Despite the usage of neural networks, recovering the landscape of 3D structures using only 2D image supervisions is still inherently ill-posed, i.e., the unknown 3D structures are of much higher dimensions than the available 2D supervisions and can easily overfit the 2D dataset. Furthermore, cryo-EM datasets often contain high-level noises. The pose parameters of each particle determined by consensus refinement present large errors since the alignment is performed between the 2D cryo-EM image with low SNR and a partial representation of the 3D structure. Without special attention, the neural network with great representation power can overfit aforementioned nuisance information in cryo-EM images by outputting unrealistic 3D structures. In this paper, in order to overcome these challenges, we adopted a new neural representation for 3D cryo-EM density map based on a convolutional architecture (9, 10), neural volumes (11). Using synthetic and real datasets, this representation has shown to be resilient to the high level of noises in 2D images and is mostly sensitive to global structural variations.

The basic assumption of 2D to 3D translation formulation induces a latent space which solely encodes the structural information from 2D images, that is, the encoder should map 2D images from the same 3D structure to the same latent code regardless of their appearances. However, the entanglement between the 3D structure and nonstructural factors in the 2D cryo-EM image poses a great challenge to implement such an encoder. Specifically, a single 3D structure can form vastly different 2D images by projecting along different angles, convoluting with different point spread functions (whose Fourier transform is contrast transfer function (CTF)) and being contaminated by noises. Therefore, an encoder as a function of 2D inputs will inevitably encode those nonstructural variations such as the projection poses, defocus parameters and noises into latent space. Another critical property of the latent space is smoothness, which guarantees that similar 3D structures are encoded by similar latent codes and alleviates the ill-posedness of 2D to 3D translation problem. Encouraging these properties of latent space can increase the amount of 2D supervisions that a 3D density map receives during training, and can improve the quality of 3D density map and the accuracy of heterogeneity analysis. In this work, by carefully designing a prior for disentangling the structural information from other factors in 2D inputs and leveraging a data augmentation pipeline, the encoder of OPUS-DSD is regularized to encode the 3D structural information in 2D images as much as possible, and the influence of projection poses and defocus parameters on latent encoding is greatly minimized.

By systematically testing on synthetic and real cryo-EM datasets, we demonstrated the superior performance of OPUS-DSD on resolving structural heterogeneity, in terms of both compositional changes and conformational dynamics, in cryo-EM datasets compared to cryoDRGN. OPUS-DSD can not only provide the necessary structural heterogeneity information to deepen our understanding about the dynamics of macromolecular systems, but also improve the resolution of highly flexible system by providing a more homogenous dataset.

## Results

### Design of OPUS-DSD

OPUS-DSD is a method for deciphering the structural heterogeneity in cryo-EM dataset using only image supervision. Overall, it contains two main parts: an encoder-decoder network to convert the 2D cryo-EM image to its corresponding 3D structure (solid red box in **Fig.1a**), and a prior in latent space that facilitates the encoding of structural information in 2D cryo-EM image (dashed red box in **Fig.1a**). The encoder-decoder network works as follows. The encoder takes a 2D cryo-EM image as input, and outputs a latent code *z* in a low dimensional space *R*^|*z*|^, while the decoder produces a 3D volume from a given latent code (**Fig.1a**). The 3D volume is represented as a discrete grid of voxels, *V*(*x*), where *x* ∈ ℝ^3^ is a point in 3D. This explicit 3D voxel grid *V*(*x*) is rendered into a 2D reconstruction with specified pose and CTF parameters according to the differentiable cryo-EM image formation model. The neural architecture of OPUS-DSD can then be trained end-to-end by reconstructing each input image and minimizing the squared reconstruction error. In order to encourage the smoothness of latent space, we adopt an instantiation of the encoder-decoder network, variational autoencoder (VAE), where *z* is assumed to be distributed as a gaussian distribution whose standard variation*σ* is parameterized by encoder (13). The latent code supplemented into decoder is sampled from the gaussian distribution of *z* given by the encoder. By forcing the decoder to reconstruct the same input using similar latent codes, the smoothness of latent space is guaranteed.

**Figure 1.**
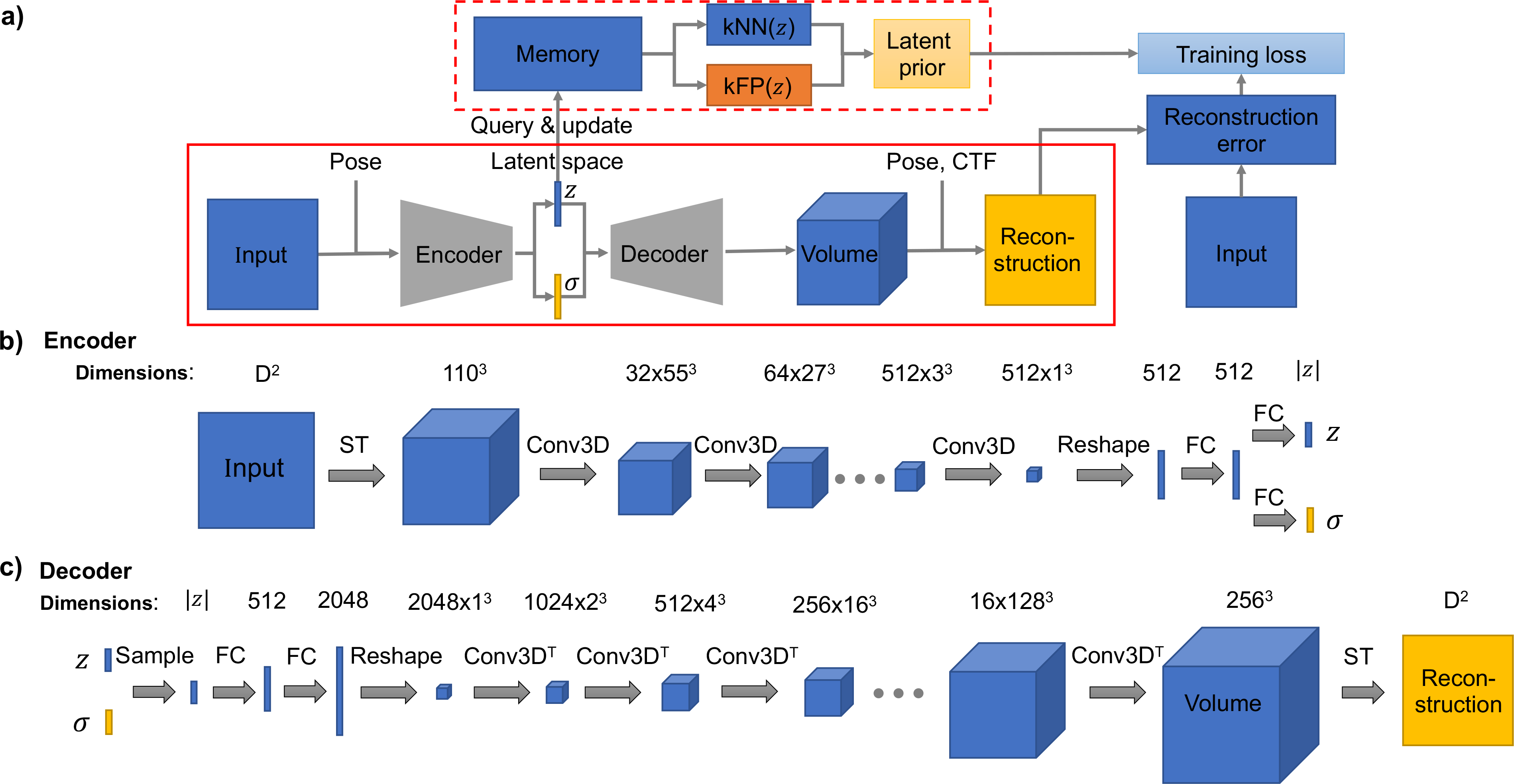
Architecture of OPUS-DSD. **a)**. Pose refers to the projection direction of input with respect to the consensus model. CTF refers to its contrast transfer function. *z* is the mean vector of latent encoding. *σ* is the standard deviation vector of latent encoding. kNN(*z*) refers to the K nearest neighbor of *z*. kFP(*z*) refers to the K farthest points of *z*. For simplicity, the standard gaussian prior for latent distribution as well as the smoothness and sparseness priors for the 3D volume are omitted in this chart. **b)**. **Architecture of the encoder of OPUS-DSD**. This diagram shows the encoder that translates a 2D cryo-EM image into the latent encoding. Top row denotes the dimensions of the intermediate tensors. The arrow links the input and output of the operation above. ST denotes the spatial transformer which backprojects the 2D image to a 3D volume. Conv3D denotes a 3D convolution. The number of channels of convolution kernel can be derived from the dimensions of its input and output. The ellipsis represents repeating the preceding operation until tensor reaches the output dimension. FC denotes the fully-connected layer. All convolution and fully-connected layers are with LeakyReLU nonlinearities with negative slope 0.2. **c)**. **Architecture of the decoder of OPUS-DSD**. This diagram shows the decoder that translates the latent encoding *z* into a reconstructed 2D projection. Conv3D^T^ denotes a 3D transposed convolution. ST denotes the spatial transformer which renders the 3D volume into the 2D image of desired resolution. All transposed convolutions except last one and fully-connected layers are with LeakyReLU nonlinearities with negative slope 0.2.

A latent prior was designed to advocate a specific type of geometry in latent space (**Fig.1a**). In the encoder-decoder architecture, resolving the structural heterogeneity amounts to encoding structural information in latent space. Namely, the variation of latent codes should mainly correlate with the variation of 3D structures. This kind of latent space induces a geometry where the similarity between 3D structures is proportional to the distance between their latent codes. In other words, the latent codes of similar 3D structures reside closely, while the latent codes of distinctive 3D structures sit at distant locations. The latent prior simultaneously collates the encodings of 2D inputs with similar 3D structures together and pushes those with different 3D structures apart, thus guaranteeing the smoothness of latent space while improving the correlation between latent codes and structural variations. For a given latent encoding, the latent encodings of 3D structures that are similar to it can be found as its K nearest neighbors (kNN), whereas the latent encodings of 3D structures that are distinct from it can be found as K furthest points (kFP) (**Fig.1a**). The latent prior is implemented by querying kNN and kFP in a dynamically updated memory bank where the latent encodings of all images are stored. The reconstruction loss guides the neural network to output an ensemble of 3D structures which fits the cryo-EM dataset, while the latent prior optimizes the geometry of the latent space to regularize the neural network to capture structural dynamics better.

The encoder network converts the 2D input to latent vector *z* by going through a series of intermediate representations which are shown as different shapes (**Fig.1b)**. The 2D projection of size *D*^2^ (square) is first converted into a pseudo 3D volume (first cube) by repeating along the *z* dimension for *D* times, and then is transformed by the inverse of its pose parameter to align with the consensus model. This transformation is performed by the spatial transformer (14). It removes the variation of 2D projection caused by the rigid transformation in 2D plane, which helps encoder to disentangle the structural heterogeneity from the pose variation. The pseudo 3D volume is down-sampled to a 1^3^ cube with 512 channels to a 512-dimension vector (first rectangle) and transform it into the latent code *z* with 512 channels using 6 consecutive strided convolutions with kernel size 4. We then flatten the 1^3^ and its standard deviation *σ* by applying two full-connected layers with non-linearities.

The 3D density maps generated by the neural network should be smooth to avoid overfitting (2, 15). The decoder adopts a convolutional architecture—neural volumes (11) to convert the latent vector to a smooth 3D volume (**Fig.1c)**. We first apply a set of fully-connected layers and non-linearities to transform the sampled latent vector (second rectangle) into a 2048-dimensional representation and reinterpret the resulting vector as a 1^3^ cube with 2048 channels. We then up-sample the 2048×1^3^ cube into a 1×256^3^ 3D volume (last cube) using 8 consecutive transposed convolutions. The kernel size of first two convolutions is 2, and the remaining convolutions are of a kernel size 4. The 2D reconstruction (square) is generated from the 256^3^ volume by the spatial transformer with given pose and CTF parameters (14).

### The performance of OPUS-DSD on synthetic NEXT complex data

NEXT complex has a dumbbell shape, where two MTR4 helicase domains located at two ends are connected by ZCCHC8 dimeric interface formed by two stranded helices (**Fig.2a**). The detailed structure determination of this complex will be reported elsewhere. In this article we will focus on testing our OPUS-DSD method and comparing it with cryoDRGN on synthetic and real cryo-EM data of NEXT complex. To synthesize data, eight different conformations are generated from the starting conformation (**Fig.2a**) by shifting the MTR4 helicase domains in relative to the ZCCHC8 center (**Fig.2b, a~h**). Each of the eight conformations has 8,000 particles. The synthetic NEXT complex dataset with 64,000 particles yielded a consensus model using cryoSPRAC (**Fig.2c**).

**Figure 2.**
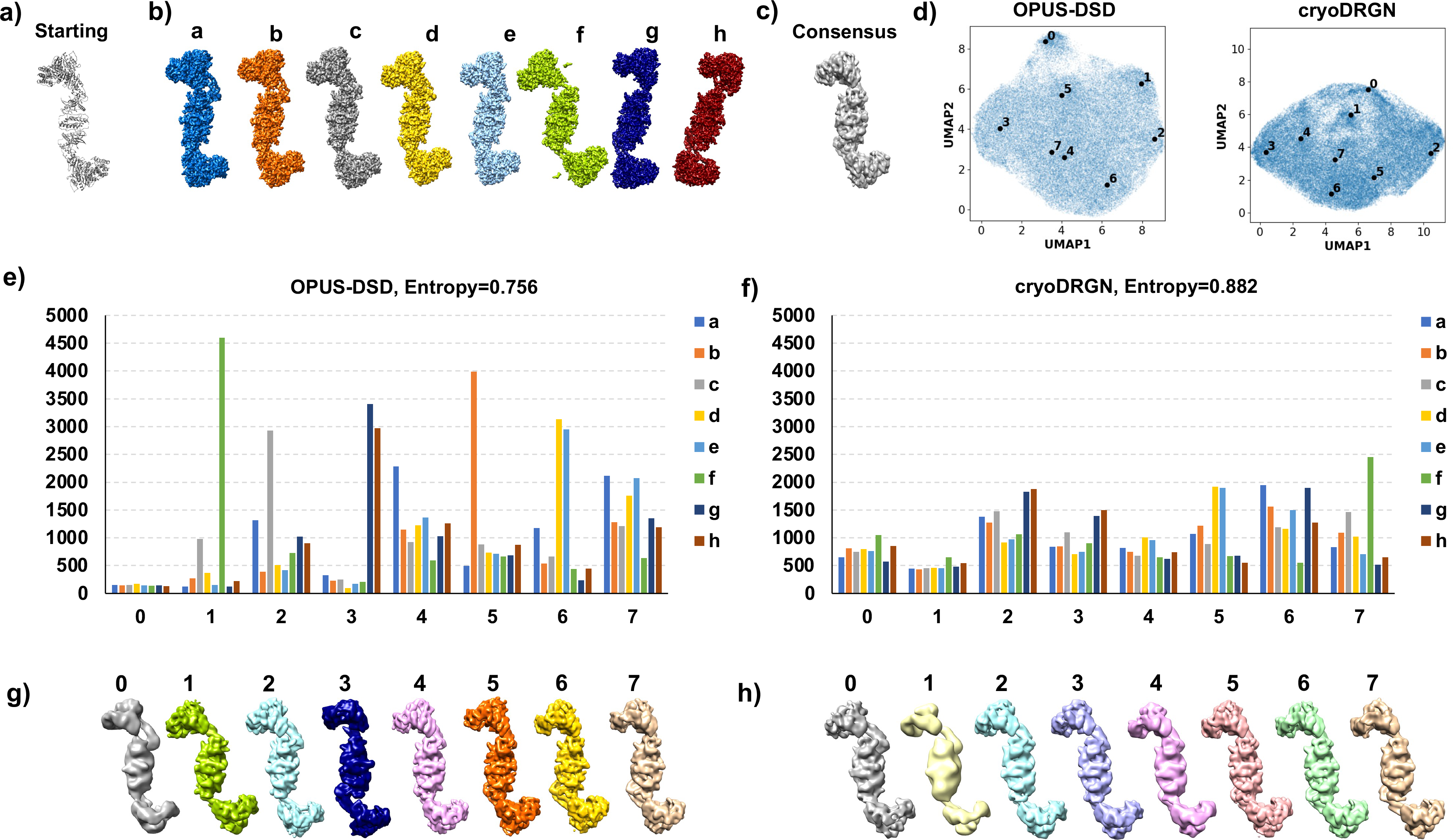
Heterogenous analyses on synthetic dataset with SNR 0.05. **a)**. The starting atomic model of NEXT complex. **b)**. The ground-truth density maps of NEXT complexes. In eight conformations, conformation *a,d,e* are similar and hard to distinguish, and conformation *f,g,h* are with larger shifts in the MTR domains. **c)**. The consensus map reconstructed by cryoSPARC. **d)**. UMAP visualizations of the 8-dimensional latent spaces of all particles encoded by OPUS-DSD and cryoDRGN. The cluster centers found by KMeans algorithm are annotated by solid black dots with corresponding label texts. **e)**. The distribution of particles in clusters in the latent space of OPUS-DSD. Bars are colored by the same profile as the ground-truth density maps in b). Entropy is defined as the average of −Σ_*i*_*p_i_* log *p_i_* of each cluster, where 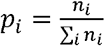, and *n_i_* is the number of particles in the ground-truth conformation *i*. **f)**. The distribution of particles in clusters in the latent space of cryoDRGN. **g)**. The 3D density maps reconstructed by particles in each class from OPUS-DSD. **h)**. The 3D density maps reconstructed by particles in each class from cryoDRGN.

When the SNR is 0.05, OPUS-DSD and cryoDRGN differ significantly in clustering results and 3D density maps (**Fig.2**). By UMAP visualizations of latent spaces, clusters are more identifiable in the latent space of OPUS-DSD compared to that of cryoDRGN (**Fig.2d**). OPUS-DSD identifies some clusters with high accuracies (**Fig.2e, Table S1**). The density maps reconstructed by particles in these classes successfully recover the ground-truth conformations (**Fig.2g**). For example, class 1 recovers the conformation *f* with about 67% of particles from conformation *f* (**Fig.2d**), and the 3D reconstruction using class 1 also resembles conformation *f* (**Fig.2g**). In contrast, cryoDRGN yields a more even distribution of the particles (**Fig.2d**) with high classification errors in all classes (**Fig.2f, Table S2**). For example, even for class 7 with the highest dominating conformation, only 28% particles are from conformation *f*. Using entropy as a metric, the average entropy of 0.756 for clusters in OPUS-DSD is much lower than that of 0.882 for cryoDRGN, which is close to the entropy of uniformly distributed conformations — 0.903. The density maps reconstructed by particles in these classes show little conformational differences except resolution changes (**Fig.2h**). The average resolution of density maps reconstructed by classes from OPUS-DSD is clearly higher than the average resolution of those from cryoDRGN, further indicating the improved classification accuracy of OPUS-DSD (**Fig.2g~h**). Judging by the quality of density map output by both methods, OPUS-DSD delivers better density maps with similar resolutions as the consensus model (**Fig.S1a**), while cryoDRGN outputs density maps with very low-resolution MTR domains and noises in the central ZCCHC8 domain (**Fig.S1b**).

When the SNR increases to 0.1 (**Fig.S2, Table S3~4**), both methods perform similarly and recover most ground-truth conformations. There are still some noticeable differences between the 3D density maps from the decoders of these two methods (**Fig.S2g~h**). The density maps from OPUS-DSD (**Fig.S2g**) have similar resolutions as the consensus model across all the complex. In contrast, the density maps from cryoDRGN overfit high-resolution information for ZCCHC8 domain in the center, while underfit MTR domains at two ends for classes with larger shifts of MTR domains (**Fig.S2h**).

In conclusion, using synthetic data, OPUS-DSD has shown to be more robust to higher levels of noise and successfully retrieves the large-scale structural variations under low SNR. On the contrary, cryoDRGN is found to be susceptible to noises and tends to overfit high-resolution information and noises that are unrelated to structural dynamics.

### The performance of OPUS-DSD on real cryo-EM data of NEXT complex

Next, OPUS-DSD is tested on experimental data of NEXT complex collected in our own group. We used two sets of data to represent different stages of single-particle reconstruction in cryo-EM. The first dataset contains 224K particles, which represents an initial stage of single-particle reconstruction after a few rounds of 2D classification. The second dataset is composed of 84K particles to represent a later stage of reconstruction where conventional classification is converged. In order to reduce the complexity of heterogeneity analysis, signals of one MTR4 helicase domain were subtracted by Relion (17).

### Test on 224K particles of NEXT complex

Consensus refinement using the 224K particles of NEXT complex reports a resolution of 5.59 Å measured by gold-standard FSC (18) (**Fig.3a**). OPUS-DSD and cryoDRGN were used to analyze the structural heterogeneity by learning a latent space with the consensus refinement result (**Fig.3a**).

**Figure 3.**
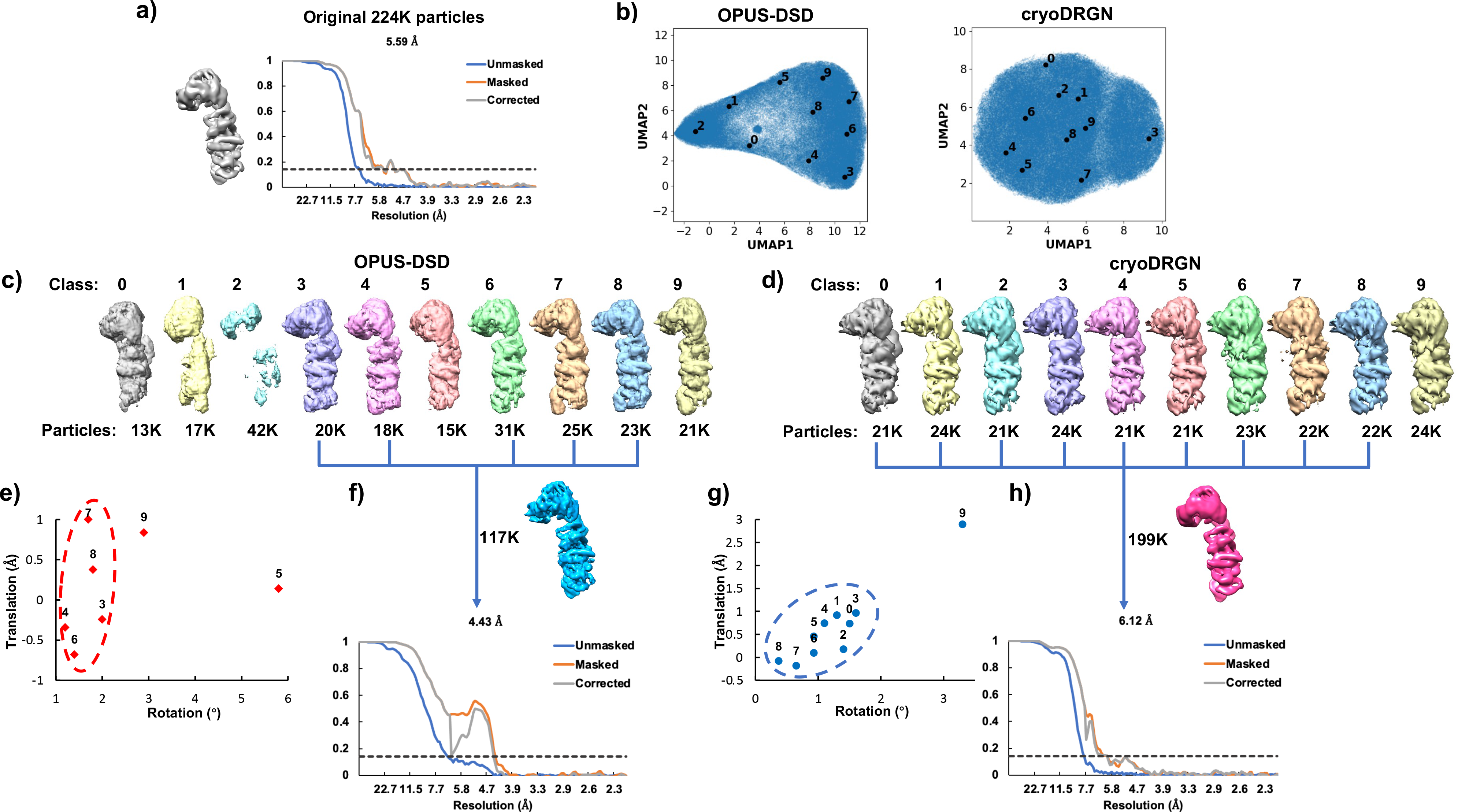
Heterogenous analyses of 224K particles of NEXT complex. **a)**. Starting consensus model and its gold-standard Fourier shell correlations (FSCs) for all particles. **b)**. UMAP visualizations of latent spaces learned by OPUS-DSD and cryoDRGN. The dots with numbers are cluster centers found by KMeans algorithm. **c)**. Ten classes reconstructed by OPUS-DSD and the corresponding number of particles in each class. There are seven classes with complete densities on the right side of UMAP (classes 3~9), and three classes with broken densities on the left of UMAP (classes 0~2). **d)**. Ten classes reconstructed by cryoDRGN and the corresponding number of particles in each class. **e)**. Translation-rotation plot for reconstructions from OPUS-DSD. The movement is measured by the scale of rigid movement of the lobe relative to the middle part of each conformation measures, which is determined by Dyndom using the consensus model as reference. **f)**. Density map and gold-standard FSCs for 117K particles by combining OPUS-DSD’s clusters with small structural variations (grouped by red circle in **e**). **g)**. Translation-rotation plot for reconstructions from cryoDRGN. **h)**. Density map and gold-standard FSCs for 199K particles by combining cryoDRGN’s clusters with small structural variations (grouped by blue circle in **g**).

For OPUS-DSD, UMAP for the latent encodings of all particles displays a bipolar distribution (**Fig.3b, left**). Classes 0, 1 and 2 on the left side of UMAP correspond to reconstructions where NEXT complex densities are not well preserved, while classes 3~9 on the right side of UMAP correspond to reconstructions where NEXT complex featured densities are complete (**Fig.3c**). In comparison, cryoDRGN on the same dataset (**Fig.3b, right**) results in reconstructions which all have complete NEXT complex densities (**Fig.3d**).

Next, the translation-rotation plots are used to visualize the structural heterogeneity for the reconstructions with the NEXT complex densities from both methods. The rotation and translation in plots refer to that of MTR helicase domain in relative to ZCCHC8 dimeric center, which are determined by Dyndom [16] where the consensus model is served as the reference (**Fig.3a**). The range of motion analysis allows us to discover clusters with relatively small structural variations for further refinement. From the translation-rotation plot for OPUS-DSD, there are five reconstructions within 2° of rotation and 1 Å of translation, forming a cluster (red circle, classes 3, 4, 6~8), and two other classes displaying a much larger extent of motion at rotation of 3° and 6° (classes 5, 9) (**Fig.3e**). For 117K particles from these five classes with a small range of motion (classes 3, 4, 6~8), non-uniform refinement (20) improves the resolution from 5.59 Å to 4.43 Å (**Fig.3f**), thus validating the compositional and conformational heterogeneity resolved by OPUS-DSD.

For cryoDRGN, there is only one reconstruction with large rotation and translation (class 9, at ~4° and ~3 Å, respectively), while the remaining nine reconstructions cluster within 2° of rotation and 1 Å of translation (**Fig.3g**). For particle from those nine classes with small motions (classes 0~8, blue circle), the non-uniform refinement yields a reconstruction of 6.12 Å, which is even worse than the resolution of consensus refinement on the original set (**Fig.3h**).

### Test on 224K particles of NEXT complex with focused refinement

The NEXT complex contains subunits presenting hugely different degrees of structural heterogeneities where the subunit can only be aligned to low-resolution in the consensus model. To further explore the applicability of OPUS-DSD, we mimicked this scenario by performing focused refinement of the ZCCHC8 dimeric region and the KOW domain of MTR4 (**Fig.4a**, the focused region is highlighted in circle). By doing so, the MTR4 helicase domain was only determined to very low-resolution, while larger displacements of MTR4 domain in relative to ZCCHC8 center were preserved in the refinement result. Thus, conformations with large MTR4 domain movements were expected to be detected.

**Figure 4.**
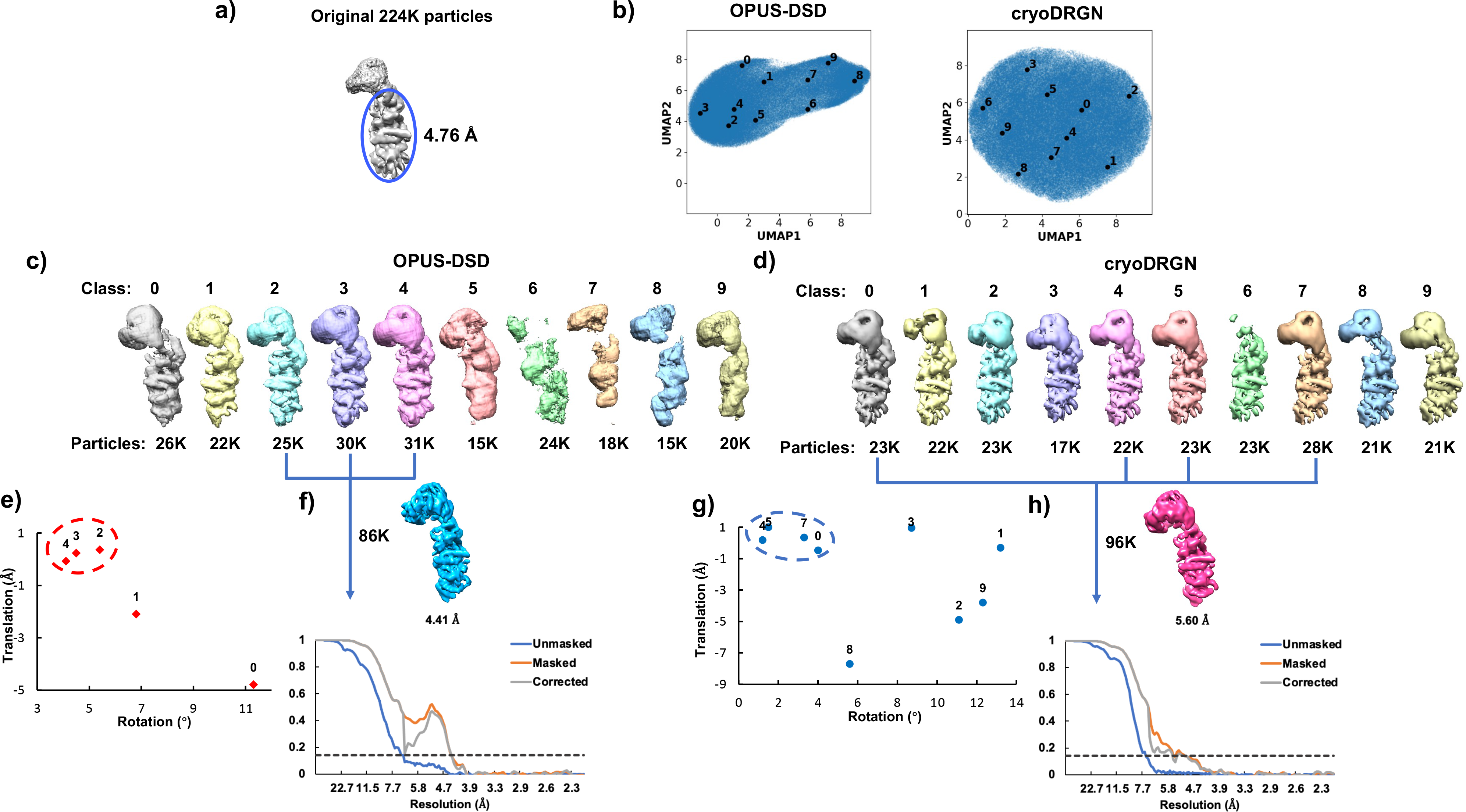
Heterogeneous analyses of 224K particles of NEXT complex with focused refinement. **a)**. Starting consensus model which was refined with a mask without the MTR4 domain using Relion. **b)**. UMAP visualizations of latent spaces learned by OPUS-DSD and cryoDRGN. The dots with numbers are cluster centers found by KMeans algorithm. **c)**. Ten classes reconstructed by OPUS-DSD and the corresponding number of particles in each class. **d)**. Ten classes reconstructed by cryoDRGN and the corresponding number of particles in each class. **e)**. Translation-rotation plot for reconstructions from OPUS-DSD. The movement is measured by the scale of rigid movement of the lobe relative to the middle part of each conformation measures, which is determined by Dyndom using the consensus model as reference. **f)**. Density map and gold-standard FSCs for 86k particles by combining OPUS-DSD’s clusters with small structural variations (grouped by red circle in **e**). **g)**. Translation-rotation plot for reconstructions from cryoDRGN. **h)**. Density map and gold-standard FSCs for 96k particles by combining cryoDRGN’s clusters with small structural variations (grouped by blue circle in **g**).

As seen from the bipolar UMAP for the latent encodings from OPUS-DSD (**Fig.4b**), classes 0~4 on the left side of UMAP corresponds to complete reconstructions, whereas classes 5~9 on the right side of UMAP are reconstructions of bad quality (**Fig.4c**). On the other hand, UMAP from cryoDRGN is circular (**Fig.4b**), and nine out of ten reconstructions show complete densities with even particle distributions (**Fig.4d**).

Translation-rotation plots from both methods (**Fig.4e~g**) show a larger range of motion than previous analysis in **Fig.3**. Classes with a small range of motion are then combined for further refinement. For OPUS-DSD, three classes clustering around rotation of 5° (classes 2~4; 86K particles) are selected (**Fig.4e**), which yield a reconstruction of 4.41 Å (**Fig.4f**). For cryoDRGN, four classes with rotation within 5° (classes 0, 4, 5 and 7; 96K particles) are selected for refinement (**Fig.4g**), and result in a reconstruction of 5.60 Å resolution (**Fig.4h**). For comparison, using the whole set of 224K particles, consensus refinement reports a resolution of 5.59 Å without focused refinement (**Fig.3a**) and 4.76 Å for the masked region (**Fig.4a**).

### Test on 84K particles of NEXT complex

We next tested both methods on a smaller dataset of 84K particles of NEXT complex, which is obtained by applying several rounds of heterogenous refinements in cryoSPARC, with the best resolution of 4.39 Å (**Fig.5a**). This scenario represents the situation where conventional classification is converged.

**Figure 5.**
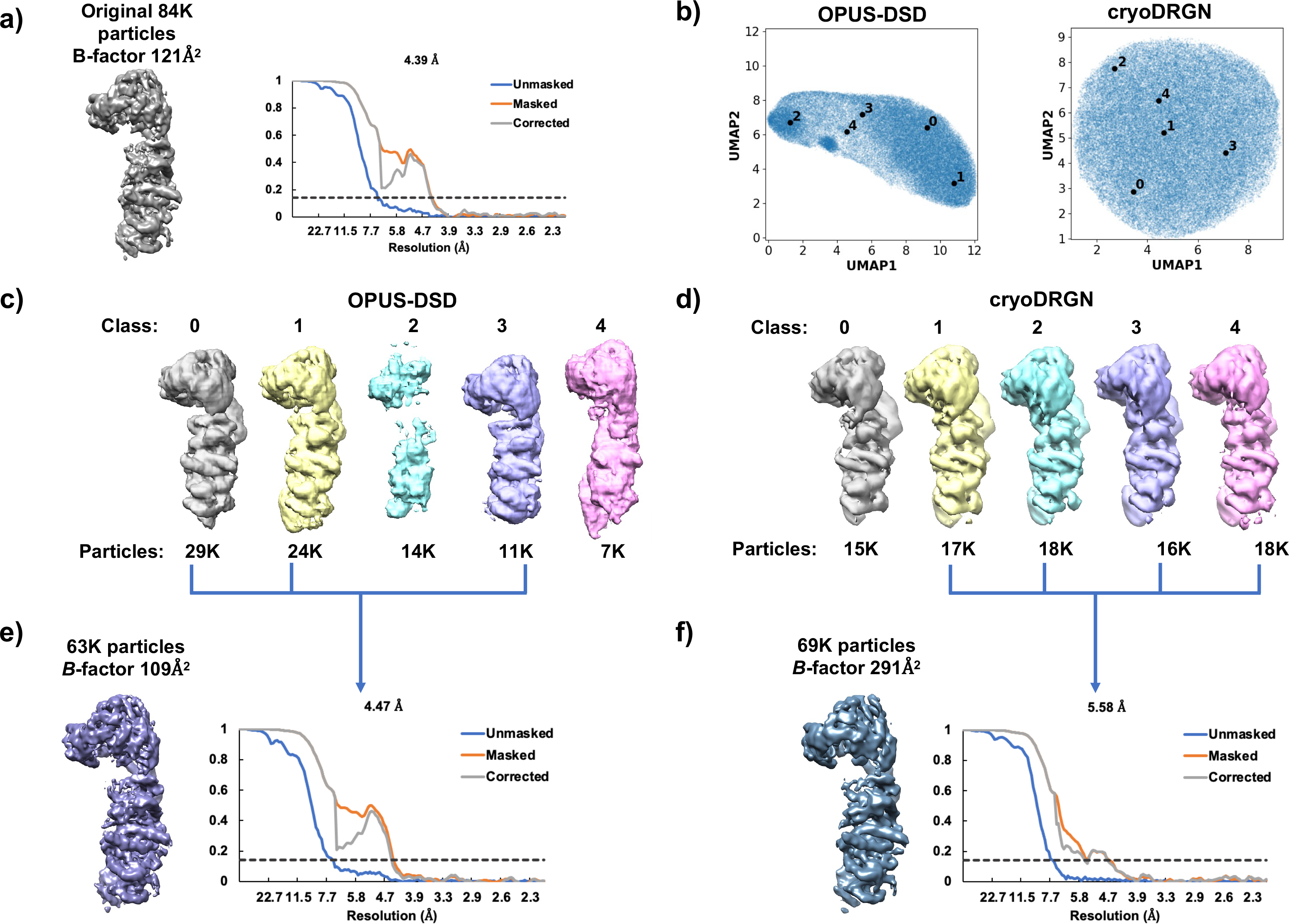
Heterogeneous analyses of 84K particles of NEXT complex. **a)**. Starting consensus model and its gold-standard FSCs for all particles. **b)**. UMAP visualizations of latent spaces learned by OPUS-DSD and cryoDRGN. The dots with numbers are cluster centers found by KMeans algorithm. **c)**. Five classes reconstructed by OPUS-DSD and the corresponding number of particles in each class. **d)**. Five classes reconstructed by cryoDRGN and the corresponding number of particles in each class. **e)**. Density map and gold-standard FSCs for 63k particles by combining OPUS-DSD’s classes 0, 1 and 3. **g)**. Density map and gold-standard FSCs for 69k particles by combining cryoDRGN’s classes 1~4.

By UMAP visualization, OPUS-DSD shows distinct conformations for particles in different locations in the latent space. Classes 0, 1 and 3 on the right side of UMAP are separated from classes 2 and 4 (**Fig.5b**). Classes 0, 1 and 3 have well preserved density of NEXT complex; in comparison, classes 2 and 4 do not show expected features of NEXT complex (**Fig.5c**). The particles in these 3 classes (63K particles in total) yield a complete reconstruction of 4.47 Å by cryoSPARC (**Fig.5e**). Although the resolution does not improve, *B*-factor decreases from 121 Å^2^ to 109 Å^2^, suggesting that the overall consistency of particles is improved (**Fig.5e**). On the other hand, UMAP from cryoDRGN results in an almost uniform distribution along two directions (**Fig.5b**). The five classes don’t distinguish each other based on UMAP nor the following reconstructions (**Fig.5b, 5d**). Non-uniform refinement on classes 1~4 (69K particles) from cryoDRGN decreases the resolution of reconstruction by 1.2 Å and increases the *B*-factor to 291 Å^2^ (**Fig.5f**). As a baseline, we determined the resolutions of four sets of particles where ~25% particles were randomly discarded. The results show that randomly discarding ~25% particles can decrease resolution by 1.2~2 Å (**Fig.S3**). This demonstrates that the clustering of these 84K particles by cryoDRGN is no better than randomly discarding a quarter of particles.

The test on the 84K particles of NEXT complex shows that OPUS-DSD could still detect real heterogeneity in the dataset at the stage where conventional classification has been converged.

### 80S Ribosome

CryoDRGN performed structural heterogeneity analysis on a large 80S ribosome dataset (EMPIAR-10028) (21) which contained 104,280 images in its original publication (8). Here OPUS-DSD is used to analyze the structural heterogeneity of a smaller dataset of 80S ribosome containing 60,363 images (EMPIAR-10002 (22)). UMAP visualization of the latent space of OPUS-DSD demonstrates good separation between the clusters (**Fig.6a**). OPUS-DSD reveals many interesting structural dynamics. First, a single 60S class is identified (class 0 in **Fig.6b**). Second, OPUS-DSD reveals the movements of an RNA strand that binds with 60S subunit (**Fig.6b**, highlighted in red dashed boxes). This movement is further demonstrated in **Fig.S4** by comparing class 9 with four other classes, where the RNA strand gradually swings from the “initial” location in class 9 to other locations. Third, the red solid boxes mark two different binding locations of an RNA strand (**Fig.6b**). Fourth, OPUS-DSD reveals the large-scale displacement of 40S subunit between classes (**Fig.6c, Fig.S5**). The density map from class 9 by OPUS-DSD is compared to the consensus model from Relion, where at the same contour level, the density map of class 9 shows more complete 40S subunit than the reconstruction by Relion using all particles due to its high flexibility (**Fig.6d**).

**Figure 6.**
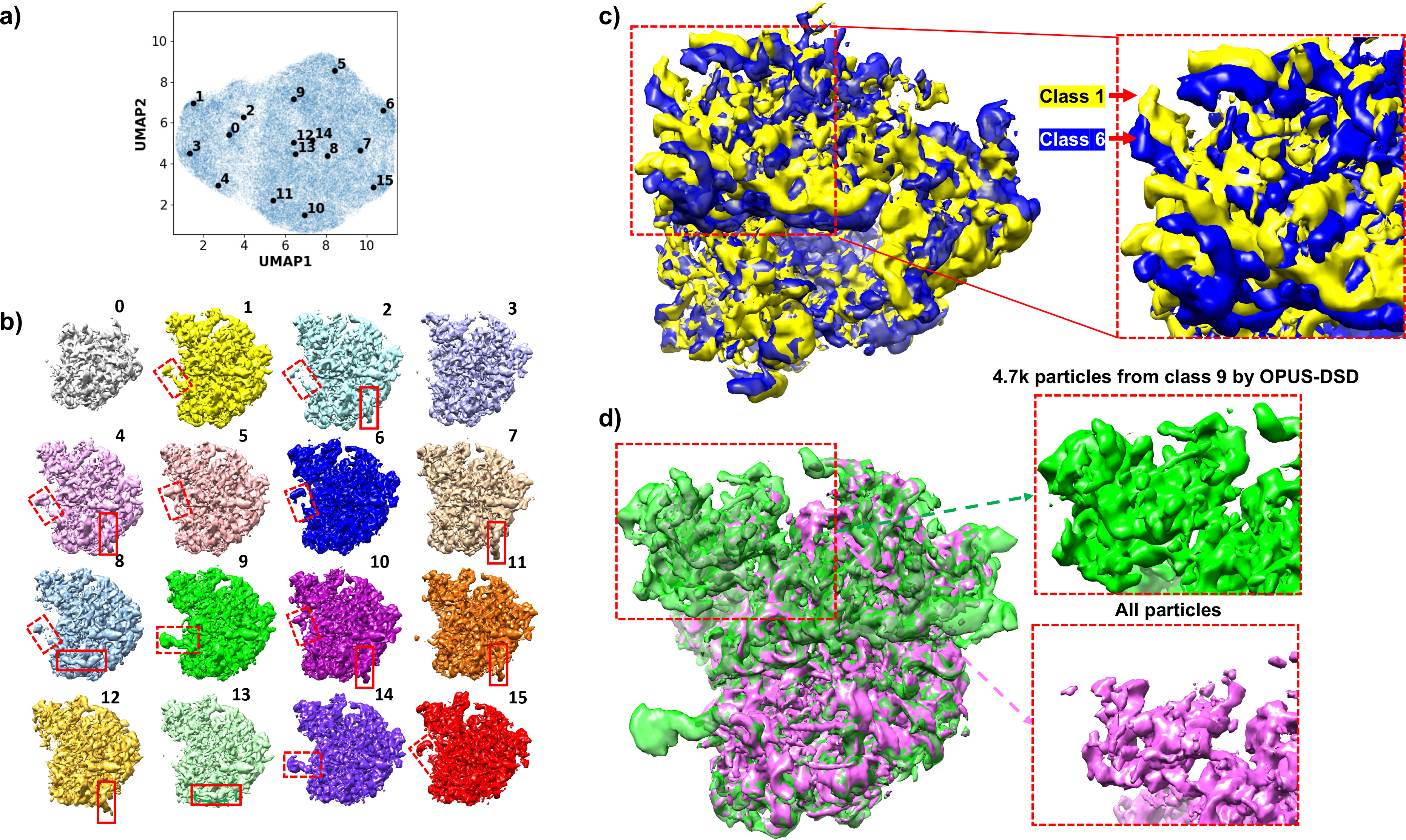
Heterogenous analyses on 80S ribosome. **a)**. UMAP visualization of the 8-dimensional latent space of all particles encoded by OPUS-DSD. Solid black dot represents the cluster center for labelled class. **b)**. Sixteen density maps reconstructed by the decoder of OPUS-DSD using KMeans clustering centers in its 8-dimensional latent space as inputs. The different locations of two RNA strands in different conformations are highlighted by red solid and dashed boxes, respectively. **c)**. Superimposition of class 1 and class 6 to show the displacement of 40S subunits highlighted in red dashed boxes. **d)**. The density map of class 9 is more complete than that reconstructed using all particles. Both maps are contoured at the same level.

## Discussion

This paper presents a new method OPUS-DSD for resolving structural heterogeneity in cryo-EM datasets. This is accomplished by learning a neural network which translates 2D cryo-EM images to 3D structures directly. The neural network is trained using only the supervision of 2D cryo-EM images whose pose parameters are determined by a consensus 3D refinement. OPUS-DSD proposed a set of methodological improvements to address the challenges in reconstructing the 3D structure from a single 2D image using neural network. The first challenge is due to the inherent ill-posedness of the 2D to 3D translation problem, since the 3D structure is of much higher dimensions and multiple 3D structures can fit the same 2D projection equally well. The ill-posedness is alleviated by encouraging the smoothness of the latent space w.r.t structural variations and the invariance of latent codes w.r.t poses and defocus. A smooth latent space guarantees that similar 3D structures are encoded by similar latent codes, while a pose invariant latent space makes the latent encodings of 2D images solely depend on their 3D structures. Therefore, the quality of 3D density map can be improved by 2D supervisions with diverse views and similar underlying 3D structures during training. Hence, OPUS-DSD proposed a latent prior to facilitate the disentanglement of structural and nonstructural factors such as projection poses and encourage encoding the structural information from 2D inputs. OPUS-DSD also leveraged data augmentation to learn a defocus invariant latent encoding. The second challenge is the low SNRs of the cryo-EM datasets. The cryo-EM images contain high level of noises and their projection poses estimated by consensus refinement often present large errors. The neural network with high representation power in cryoDRGN can easily overfit nuisance information in images which is unrelated to the structural variations. The neural network in cryoDRGN also relies on an explicit pose encoding to model the 2D projection. This approach may be susceptible to the pose assignment errors. These features of cryoDRGN yield unsatisfactory 3D density maps on datasets with low SNRs. The systematic errors in 3D density maps will propagate into latent space during training and ultimately lead to noninformative latent encodings, which yields spurious structural heterogeneity resolving results. These architectural drawbacks in cryoDRGN are demonstrated using both synthetic and real datasets in our study here. On the other hand, OPUS-DSD overcomes these peculiarities by adopting a 3D convolutional architecture [10]. Instead of representing the 2D projection using explicit pose encoding, OPUS-DSD reconstructs the 2D projection by directly projecting a 3D volume. This convolutional architecture appears to be robust to high level of noises and incorrect pose assignments in 2D cryo-EM images. It also focuses on capturing large-scale structural variations while being resilient to the irrelevant high-resolution structural information and noises. An advantage of this approach is that it allows one to further improve the fidelity of the 3D density maps by using the traditional sparseness and smoothness regularizers such as total variation and LASSO (15, 23, 24). The smooth 3D density maps produced by the decoder reduce modelling errors, generate an informative latent space and greatly increase the resolving power of structural heterogeneity. Moreover, OPUS-DSD employs a validation set to objectively monitoring the neural network training process, thus further reducing the problem of overfitting nonstructural heterogeneity.

OPUS-DSD was applied to different stages of cryo-EM data processing: 1) dataset at an early stage with a large amount of particle heterogeneity; 2) when classification is converged. For stage 1), conventional methods may need much more rounds of classification before a high-resolution reconstruction can be obtained, while OPUS-DSD could obtain high-resolution reconstruction and reveal structural heterogeneity in just one round. For stage 2), OPUS-DSD could be used to further detect particle heterogeneity. By purifying the dataset further, the reconstruction could be further improved. OPUS-DSD should find ample applications to cryo-EM data with local high flexibilities that are not rigid-body movements. In such cases, rigid-body analysis such as multibody refinement (6) would be less efficient, which requires further dividing flexible regions into smaller units to fit the movement, and different division schemes for different modes of movements. For example, the rotation of 40S subunit and the swing of RNA strand in 80S ribosome should be represented by two different division schemes to resolve them when using multibody refinement. In contrast, OPUS-DSD requires no prior knowledge about the underlying dynamics and can automatically discern different states from inputs. When the SNR of a dataset is sufficient, OPUS-DSD can even recover densities that are missing in conventional reconstruction as shown in **Fig.6d** for 80S ribosome. More importantly, OPUS-DSD has the advantage of exploring conformational and compositional changes in a unified framework.

In conclusion, OPUS-DSD is a robust, accurate and versatile tool for resolving structural variations. It can not only deepen our understandings about the dynamics of biological systems but also facilitate structure determination by providing more homogenous datasets to achieve higher resolutions.

## Methods and Materials

### Notation

The notations used in this paper are listed as following. *V* often represents the 3D structure. *X* represents the 2D projection of the 3D structure. For a vector *x* ∈ ℝ^*N*^, 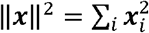 is the sum of square of the vector 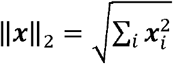 represents the *l*_2_ norm of the vector *x*. ||*x*||_1_ = ∑_*i*_|*x*_*i*_| represents the *l*_1_ norm of the vector *x*. ·^*T*^ represents the transpose of a matrix. Var represents the variance of a random variable. *SO*(3) represents the 3D rotation group.

### Image formation model

The image formation model of cryo-EM is typically defined in frequency domain. Since the images collected by cryo-EM are 2D projections of a 3D molecular structure, the Fourier transform of the image has the following relation with the Fourier transform of the 3D molecular structure according to projection-slice theorem. Let the 3D molecular structure be *V*, and its Fourier transform be 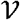. Assume an *N*^2^ image *X* is formed by rotating a *N*^3^ 3D volume *V* with the Euler angle set *ϕ* and projecting along with the *z* axis, using projection-slice theorem, the Fourier transform of the image 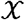 can be expressed as

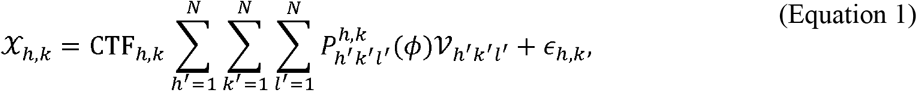

where 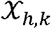 is a component of the Fourier transform of the image *X* whose spatial frequency vector is [*h*,*k*], CTF_*h*,*k*_ is a component of the contrast transfer function (CTF) (25), 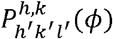 is the slice operator which cuts out a plane in the 3D Fourier transform 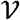 according to the Euler angle set *ϕ* and *ϵ*_*h,k*_ is the noise that corrupts the projection. The 2D image *X* can further undergo a set of 2D rigid transformations such as rotation and translation in projection plane. Since those transformations can be easily corrected using determined pose parameters, they will be ignored in our discussions. We elaborate on the slice operator *P^ϕ^* by giving its formal definition. Let the index of a voxel in the 3D Foruier transform 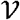 be [*h*′, *k*′, *l*′] and the index of the corresponding pixel of the Fourier transform of the image is [*h*,*k*] the slice operator *P^ϕ^* transforms the 3D index to 2D index by the following equation,

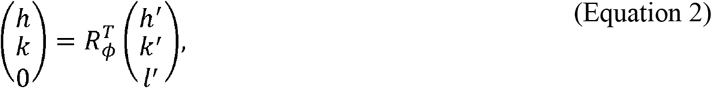

where *R_ϕ_* is a rotation matrix parameterized by Euler angle set *ϕ*. The counterpart of the slice operator in real space is the projection operator. The projection operator rotates a reference volume according to a Euler angle set. Let *V* be the template volume, and *V*′ be the volume rotated by the Euler angle set *ϕ*, the projection operator rotates the 3D template volume by the following equation,

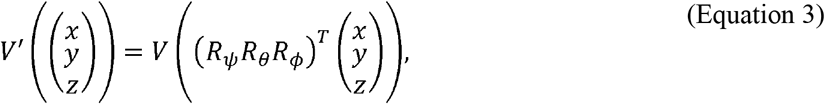

The projection operator then generates the 2D projection *X* by summing along the *z* axis of the rotated volume *V*′,

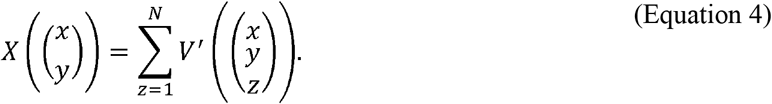

In summary, the 3D volume undergoes a rotation in 3D and 2D rigid transformations before transforming to a 2D projection. OPUS-DSD represents structure by 3D volume in real space. The 2D cryo-EM reconstruction is obtained as follows in it; the 3D volume is rotated by the spatial transformer (14), the rotated 3D volume is projected along *z* axis to form a 2D projection, and the 2D projection further undergoes defocus correction to generate the 2D cryo-EM reconstruction by applying contrast transfer function as in **Error! Reference source not found**. . The 2D cryo-EM image *X* can be expressed as a function of the 3D volume *V*, the projection angle *P* and the defocus parameters of microscope *u* which defines the CTF, i.e., *X*(*V,P,u*). Therefore, the whole image formation process is differentiable and is suitable for end-to-end training.

### cryoDRGN

CryoDRGN proposed a novel representation to model the 2D projection from 3D volume. The projection can be described as a continuous function *V*:ℝ^3^ → ℝ. CryoDRGN approximates the function *V* directly by using multilayer perceptron (MLP). Instead of supplementing the 3D coordinates into the MLP, cryoDRGN encodes the coordinates using a specific positional encoding (8). The 2D projection along a specific angle of the 3D volume *V* is computed by supplementing positional encodings of the coordinates of its corresponding slice (Equation 2) pixel by pixel. Different structures are generated by concatenating latent encoding to the positional encodings. CryoDRGN is trained using *β*-VAE (27).

### Structural Disentanglement Prior

A main component of OPUS-DSD is a novel prior in latent space which facilitates the disentanglement between the 3D structural heterogeneity and nonstructural information and encoding the 3D structural heterogeneity. In the framework of encode-decoder network, resolving the structural heterogeneity in cryo-EM dataset can be formulated as learning a latent space that only captures the 3D structural information. Each latent code represents a unique 3D structure in this space, ***z***~*V*. Given a latent code, the decoder network can produce the corresponding 3D volume *g*(*V*|***z***). The similarity between 3D structures correlates with the distance between their latent codes. From the perspective of decoder, the latent space can be encouraged to encode only the structural information by explicitly supplementing the nonstructural information like poses and defocus parameters into the reconstruction of 2D projection from 3D volume. As long as the nonstructural information is accurately modelled, nonstructural information in 2D images will stop propagating from the decoder into the latent space during backpropagation. However, when the pose assignments of 2D images are erroneous, the 3D reconstruction from the decoder will be distorted to account pose assignment error. Therefore, this approach is greatly affected by the accuracy of pose assignment. The structural heterogeneity power of the neural network will quickly deteriorate as the SNR of dataset drops. The disentanglement of structural contents from pose parameters in cryoDRGN is achieved via this approach (28). From the perspective of encoder, learning a structural latent space equivalents to learning a projection and defocus invariant encoding for all 2D cryo-EM images from the same 3D structure, namely, *f*(*X*(*V*, *P*, *u*)) = *f*(*X*(*V*, *P*′, *u*′)), ∀*P*, *P*′ ∈ *SO*(3), ∀*u*, *u*′ ∈ ℝ^*n*^. The 2D cryo-EM image is a function of 3D structure *V*, projection angle *P* and defocus parameters *u, X*(*V*, *P*, *u*). It can vary considerably by changing the projection angle and defocus parameters while fixing the underlying 3D structure, namely *X*(*V*, *P*, *u*) ≠ *X*(*V*, *P*′, *u*′) as long as *P* ≠ *P*′ or *u* ≠ *u*′. In order to approximate such a latent space which primarily encodes 3D structural information, the encoder network should disentangle the 3D structural variations in 2D inputs from other factors such as projection angle and defocus parameters. Thus, the encoder network in OPUS-DSD is encouraged to generate latent encodings with larger variations for 3D structural changes in 2D inputs, while being relatively insensitive to the poses and contrast changes, namely, Var(*∂f(X(V, P, u))/∂V*) ≫ Var(*∂f(X(V, P, u))/∂Δ*),Δ = *P* or *u*, where variances are taken over all images. Specifically, the disentanglement is achieved by adding attracting forces and expelling forces between the encodings of specific combination of images. The attracting forces restrain the distance between the latent codes of images from similar 3D structures, while the expelling forces encourage the separation between the latent codes of different 3D structures. Firstly, the 3D structural heterogeneity is most discernible in 2D cryo-EM images which are of the same projection angle, namely, the variation of *X*(*V, P, u*) can be mainly attributed to *V* when conditioned on *P*. The expelling force is added between the latent codes of those pairs of cryo-EM images to amplify the difference between latent codes for images from different 3D structures, which encourages the encoder to discern images from different structures. Next, to encourage the smoothness of latent codes, the distances of the latent codes of similar images are restrained to prevent over separation. To formally define the prior, since the projection angles form an *SO*(3) group which can be discretized into a number of classes using HEALPix (29), we classify 2D images into different projection classes according to their projection angles. Suppose the encoder network is *f*, the projection class of image *X*_*i*_ is *P*_*i*_, the projection class of image *X*_*j*_ is *P*_*j*_ inspired by the objective function of UAMP (16), the prior for the encoder network which encourages the encoding of structural information for images within the same projection class *P_i_* can be expressed as

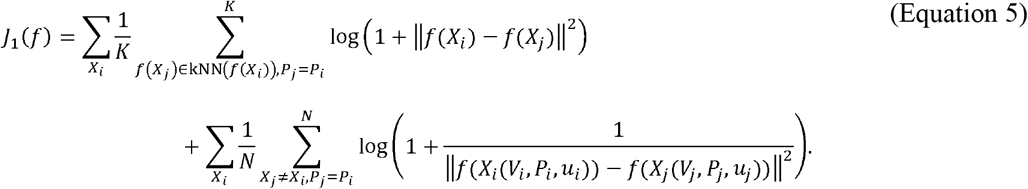

where kNN refers to the *k* nearest neighbors of the image measured by Euclidean distance in latent space *K* refers to the number of nearest neighbors, and the second summation in the second term is over all images within the same class as image *X*_*i*_ except itself, *N* refers to the size of the projection class of image *X_i_*. For images with similar encodings, the attracting prior reaches minimum when *f*(*X_i_*) = *f(X_j_)*. In contrast, the minimum of expelling prior is obtained at ||*f(X_i_)* − *f(X_j_)*|| → ∞ thus the encoder is forced to amplify the structural differences between *X*_*i*_ and *X*_*j*_. The second part of our prior encourage learning pose invariant latent encodings. This is achieved by collating the latent codes of different projections of the same 3D structure together. The projections with similar latent codes are considered to be from the same structure. The attracting force can be added among them. To encourage the separation of different 3D structures, a repulsion prior is added to an image and its *k* farthest point in latent space. Formally, let the *k* nearest neighbors of the image *X*_*i*_ in latent space be kNN(*f*(*X_i_*)), the projection class of image *X_i_* be *P_i_*, the *k* farthest point of the image *X_i_* in latent space be kFP(*f*(*X_i_*)), our prior for encouraging the projection pose invariance of latent encoding can be expressed as,

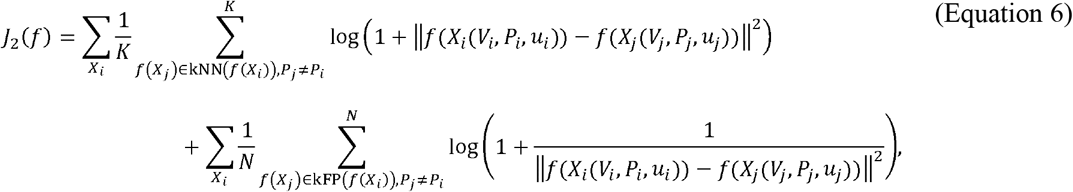

where the second summation in both terms are over all images which don’t belong to the same projection class as *X*_*i*_, *K* refers the number of nearest neighbors of *X*_*i*_ and *N* refers to the number of farthest points of *X_i_*, which are both tunable hyperparameters. The whole prior for structural disentanglement is the sum of (Equation 5) and (Equation 6), which can be written as

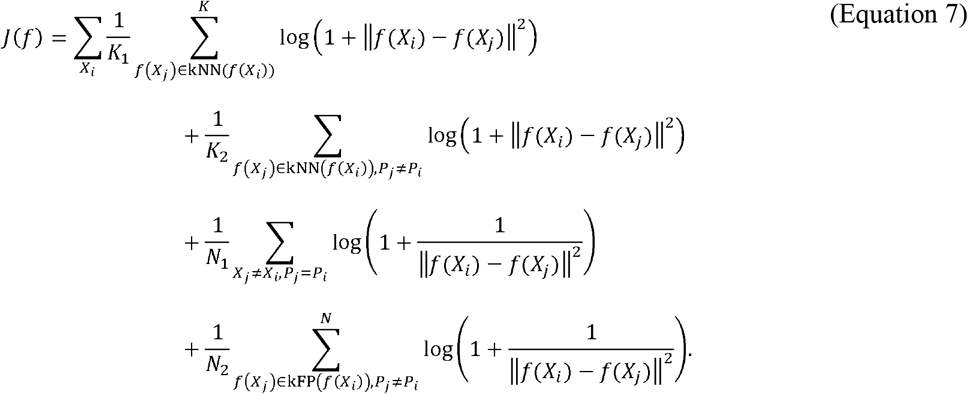

The remaining problem is to design an efficient method to compute the structural disentanglement latent prior. (Equation 5) can be approximated using a batch of images from the same projection class. For (Equation 6), since the cryo-EM dataset of contains hundreds of thousands of images, it is computationally inefficient to compute kNN and kFP for each image over all images. We compute them approximately for each query by using a subset of images from projection classes other than the query’s projection class. This subset is randomly sampled from a memory bank that stores all latent encodings. The distances between the query latent code and samples are then computed and sorted ascendingly. The first *K*_2_ latent codes with shortest distances are designated as the nearest neighbors of the query. The last *N*_2_ latent codes with longest distances are denoted as the farthest points of the query.

### Data Augmentation

The final ingredient of OPUS-DSD to encourage a defocus invariant latent space is a data augmentation pipeline. The contrast transfer function blurs the cryo-EM 2D projection while strongly modulating its contrast [20]. To make the network to be invariant to the defocus variation, the network uses input image with its contrast and brightness randomly adjusted to reconstruct the original image. This data augmentation pipeline increases the robustness of encoder network to the contrast variations of inputs, and reduces the impact of defocus parameters on latent encodings.

### Training objective

The smoothness of latent space is of paramount importance to the generalizability of the generative model. To generate plausible samples during a traversal through the latent space, the generative model needs to smoothly interpolate between training samples. One way to achieve the smooth interpolation between samples is using variational autoencoder (13). Instead of producing a deterministic latent code, the encoder approximates a posterior distribution of latent code conditioned on a image *X*, *f*(***z***|*X*), which is often assumed to be a diagonal gaussian distribution for simplicity. The encoder parameterizes this diagonal gaussian distribution by outputting its mean and variance. The smoothness of latent space is encouraged by increasing the overlapping between the approximate posterior distributions of images, namely, restraining the deviation of approximate posterior distributions from the standard gaussian distribution. Furthermore, to generate plausible 3D volume, total variation and LASSO priors are imposed on the reconstructed 3D volume to encourage its smoothness and sparseness (23, 24). Using VAE, let *f* be the encoder network, *g* be the decoder network, *F* represent the image formation process given the 3D volume *V*, projection angle *θ*, and defocus parameters *u*, the dimension of the 3D volume be *N*^3^, the objective function for learning a neural network to resolving 3D structural heterogeneity can be expressed as,

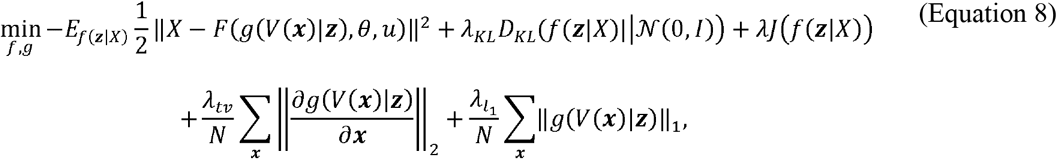

where the first term is the expectation of error between the ground-truth image *X* and the reconstructions *F*(*g*(***z***),*θ*,*u*) over the distribution of latent code 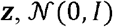 is the standard gaussian distribution, *D_KL_* is the Kullback-Leibler (KL) divergence between the distribution given by encoder network *f* and the standard gaussian 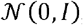, the term *J*(*f*(***z***|*X*)) is the structural disentanglement prior to encourage the encoding of 3D structural information in latent space, 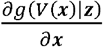 denotes the gradient of 3D volume at grid point ***x***. Computing the expectation in (Equation 8) is intractable, the VAE uses reparameterization trick to approximate the expectation integral (13).

### Training

Our network is trained using only 2D image supervision. The projection pose parameters of 2D images are determined by consensus 3D refinement in Relion or Cryosparc (5, 30). The 2D images are randomly split into a training set and a validation set using a 9-1 split. The images in validation set are not involved in training neural network and used to evaluate the reconstruction quality of the neural network only. The computation of structural disentanglement prior requires us to perform intra-projection class and inter-projection class comparisons. Hence, the training images are classified into 48 different projection classes according to their first two Euler angles using HEALPix (29) before supplementing to training. At each iteration, a projection class is selected uniformly, and a batch of images within the selected projection class is sampled by a customized batch sampler. The structural disengtanglement prior can be easily computed using such a batching process. The image sample is passed to the encoder which outputs a sample of the latent distribution *f*(***z***|*X*). The decoder reconstructs a 3D volume according to the latent code. Total variation and LASSO priors are imposed on the reconstructed 3D volume to encourage its smoothness and sparseness (23, 24). The 3D volume is rendered into a 2D reconstruction using the projection angle of the input according to the cryo-EM image formation model. The inputs and reconstructions together with the latent codes are used to construct the training loss as in (Equation 8). The network is trained by minimizing the training loss using the Adam optimizer (31).

### Implementation

The neural network of OPUS-DSD is implemented with Pytorch with automatic differentiation support (32). Using 4 Nvidia V100 GPUs, the training of 1000 images takes around 1 minute.

### Hyperparameter settings

In our experiments, we set the restraint strength for KL divergence *λ_KL_* = 0.01, the restraint strength for the total variation of 3D volume *λ_tv_* = 1 and the restraint strength for LASSO 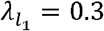. The restraint strength for structural disentanglement prior was set according to the level of signal in the cryo-EM datasets. For 80S ribosome with high level of signal, we set *λ* = 1. For synthetic data and real NEXT complex datasets, we set *λ* = 0.5. Overall restraint strength was further weighted by a cyclical annealing schedule to avoid posterior collapse (33). The learning rate was set to 10^−4^ and decayed by 0.95 at each epoch for all experiments. The batch size was set to 72 during training. All trainings were performed on 4 Nvidia V100 GPUs. The number of samples for interclass comparison is set to 12800. The number of kNN for interclass comparison is set to 128, while the number of kFP for interclass comparison is set to 512. The number of kNN for intraclass comparison is set to 4. The latent encodings of particles are updated with momentum to be 0.7.

### Experimental Protocol

For each system, the structural heterogeneity analyses were performed on the same consensus refinement result for different methods. In all experiments of this paper, the dimension of latent space is fixed to be 8 for all methods. For cryoDRGN, due to the lack of validation metric, the number of training epochs for the model chosen for result analysis was judged by the quality of density maps reconstructed by corresponding model. However, OPUS-DSD selected models for further analyses according to the reconstruction error on the validation set. The structural heterogeneity was revealed by performing clustering in latent space which encodes the structural variations. The 3D density maps reconstructed from the centroids of clusters can be designated as the representative conformations. The informative latent space with well separated clusters should be easily clustered by the simple KMeans clustering algorithm (34). Hence, throughout this paper, the structural heterogeneity analyses performed by different methods were evaluated by comparing the KMeans clustering results over the latent encodings of 2D images. The latent spaces of different methods were also visualized in 2D by UMAP (16). Informative latent space usually forms distinct clusters in UMAP visualization.

### Synthetic Data Preparation

The synthetic dataset comprised of 64,000 2D images of 8 different conformations. Eight different conformations were generated from the starting conformation by shifting the MTR4 helicase domains in relative to the ZCCHC8 center. The density maps for these 8 conformations were generated by *molmap* command in Chimera from corresponding atomic models (35). The 2D projection is obtained by projecting the density map of a conformation into 2D plane along a randomly sampled projection angle. The 2D projection was further convolved with randomly sampled contrast transfer function and contaminated by specific level of noises to generate the final synthetic 2D image. Each conformation produced 8,000 synthetic images in such a way.

## Acknowledgements

This work was partially supported by Shanghai Municipal Science and Technology Major Project (Award Number 2018SHZDZX01) and ZJLab. This work was also supported by National Key Research and Development Program of China (Award Number 2021YFF1200400).

## Supplementary Figure legends

**Figure S1. Reconstructions of OPUS-DSD and cryoDRGN on synthetic dataset with SNR 0.05. a)**. 3D density maps reconstructed by the decoder of OPUS-DSD using cluster centers. **b)**. 3D density maps reconstructed the decoder of cryoDRGN using cluster centers.

**Figure S2. Heterogenous analyses on synthetic dataset with SNR 0.1. a)**. The starting atomic model of NEXT complex. **b)**. The 8 ground-truth density maps of NEXT complexes. **c)**. The consensus map reconstructed by cryoSPARC. **d)**. UMAP visualizations of the 8-dimensional latent spaces of all particles encoded by OPUS-DSD and cryoDRGN. **e)**. The distribution of particles in clusters in the latent space of OPUS-DSD. Bars are colored by the same profile as the ground-truth density maps in b). Entropy is defined as the average of −Σ_*i*_*p_i_* log *p_i_*, of each cluster, where 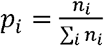, and *n_i_* is the number of particles in the ground-truth conformation *i*. **f)**. The distribution of particles in clusters in the latent space of cryoDRGN. **g)**. The 3D density maps reconstructed by particles in each class from OPUS-DSD. **h)**. The D density maps reconstructed by particles in each class from cryoDRGN.

**Figure S3. Reconstructions by cryoSPARC non-uniform refinement after randomly discarding a quarter of original 84K particles in four parallel trials**. The resolutions of models are in range of 5.60 Å to 6.35 Å.

**Figure S4. The movement of an RNA strand in 80S ribosome revealed by OPUS-DSD**. The reference density map of class 9 is colored in green and semi-transparent. The density maps shown in sequence are classes 8, 4, 6, and 5, and have increasing larger displacements in the RNA w.r.t class 9 marked by red dashed boxes.

**Figure S5. The movement of 40S subunit in 80S ribosome revealed by OPUS-DSD**. The reference density map is colored in violet and semi-transparent. The density maps shown in sequence are classes 2, 11, 13, 8, 7, and 15, whose locations in UMAP visualization vary from left to right according to UMAP1 and have increasingly larger displacements in 40S subunit w.r.t class 4 marked by red dashed boxes.

