## Supplementary Figures for "OPUS-DSD: Deep Structural Disentanglement for cryo-EM Single Particle Analysis"

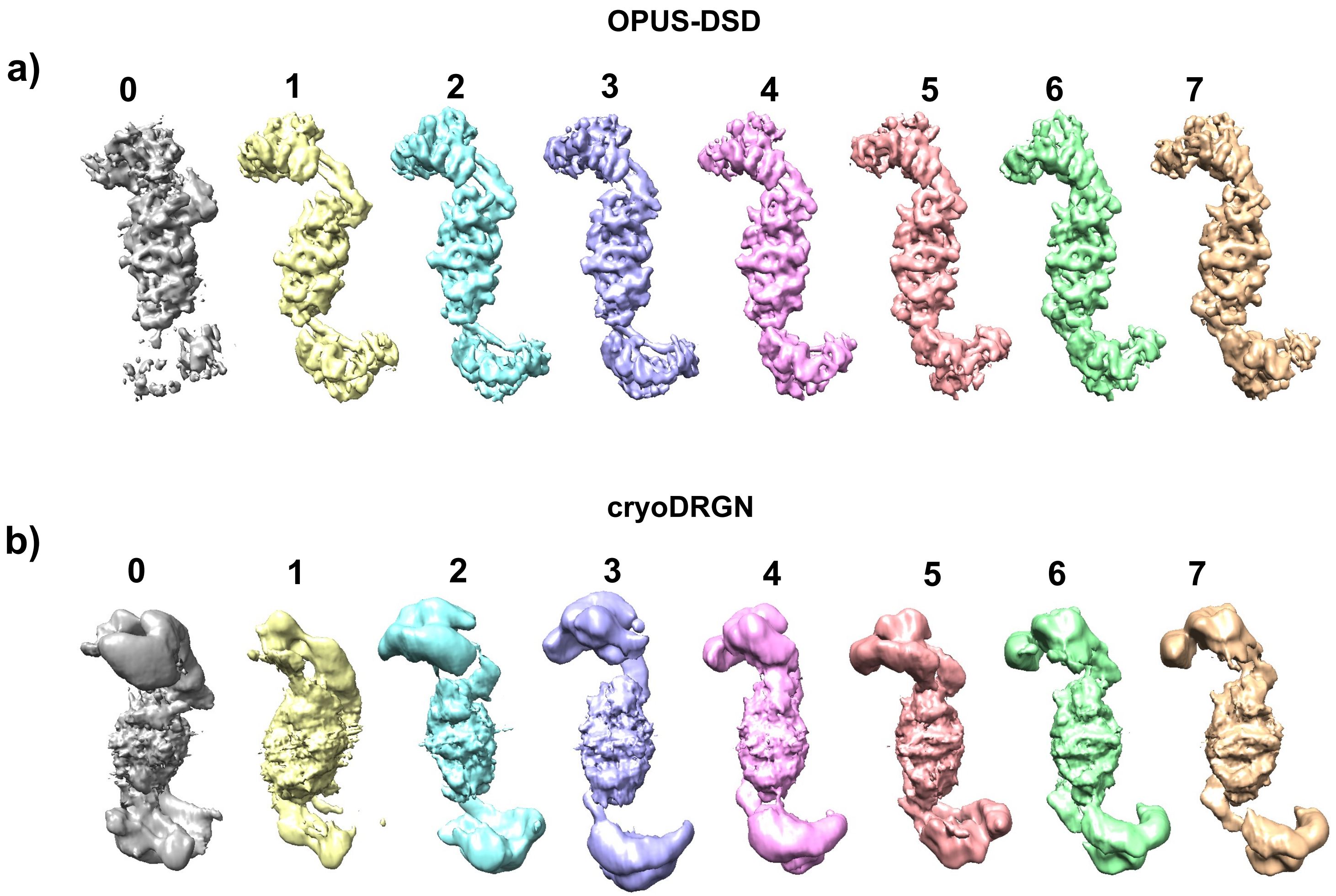


Fig.S1

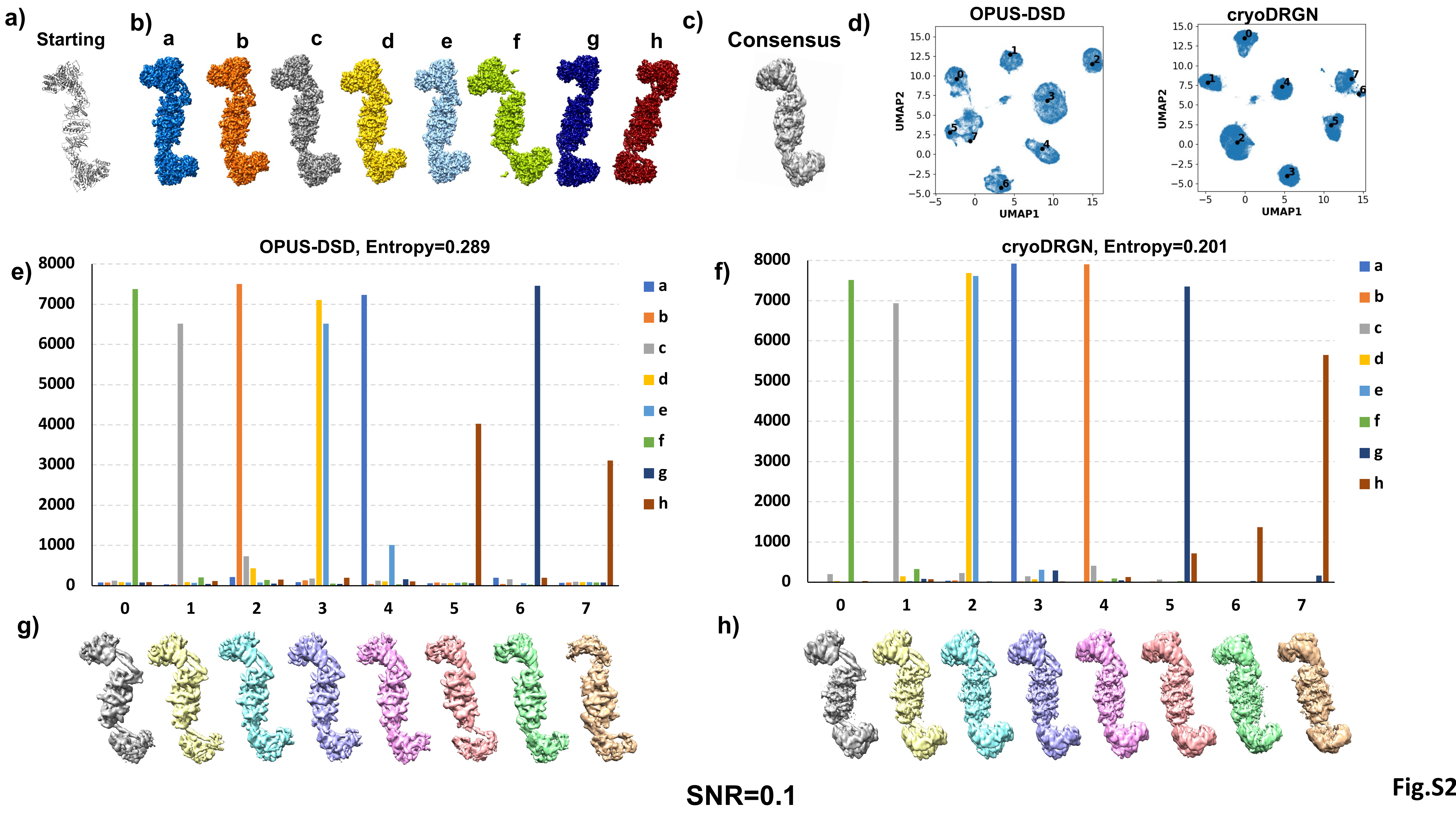

a)
Starting
b)
a
b
c
d
e
f
g
h
c)
Consensus
d)
OPUS-DSD
cryoDRGN
UMAP1
UMAP2
e)
OPUS-DSD, Entropy=0.289
f)
cryoDRGN, Entropy=0.201
g)
h)
SNR=0.1


Fig.S2

**Refinements after randomly discarding quarter of original 84k particles**

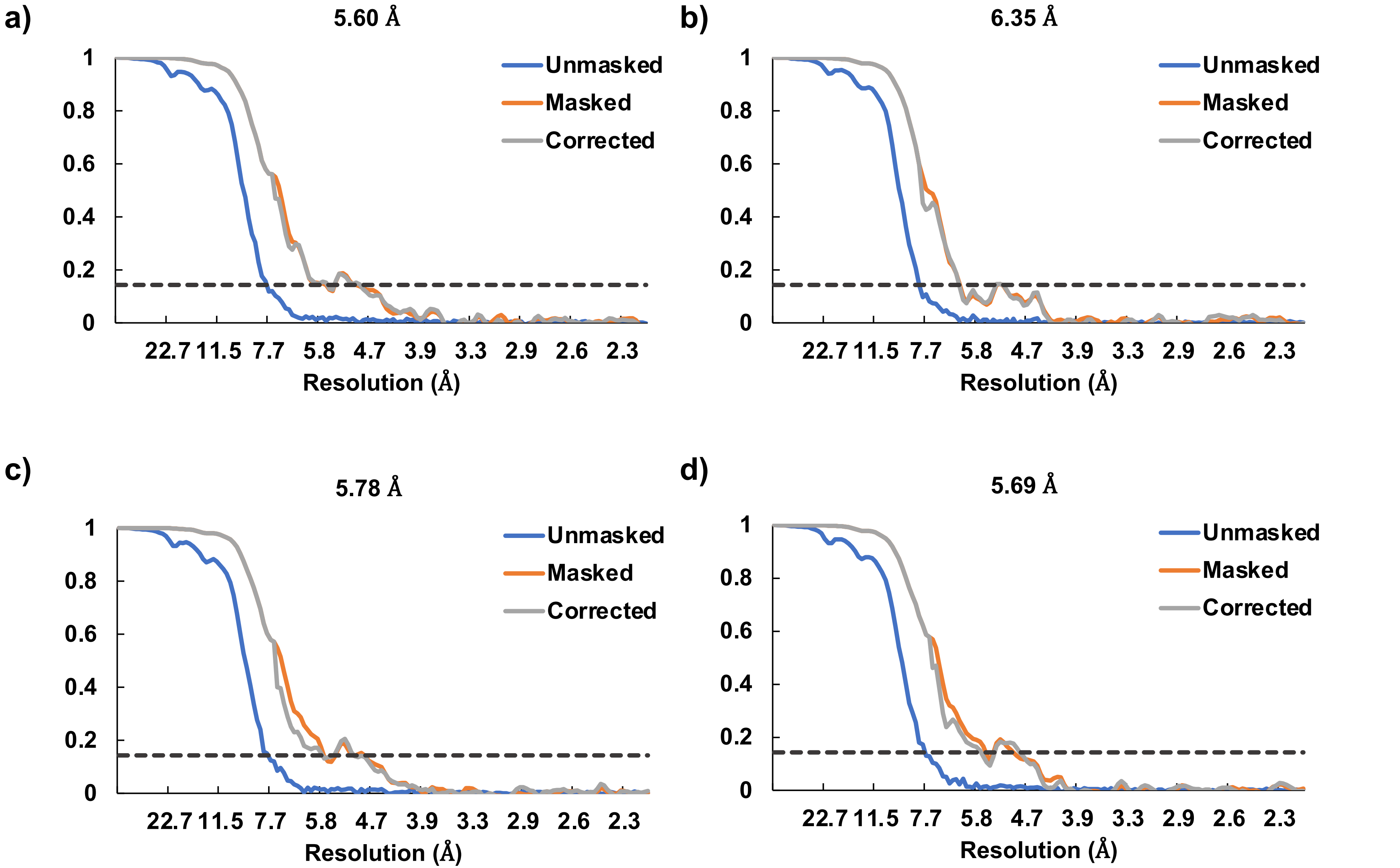


Fig.S3

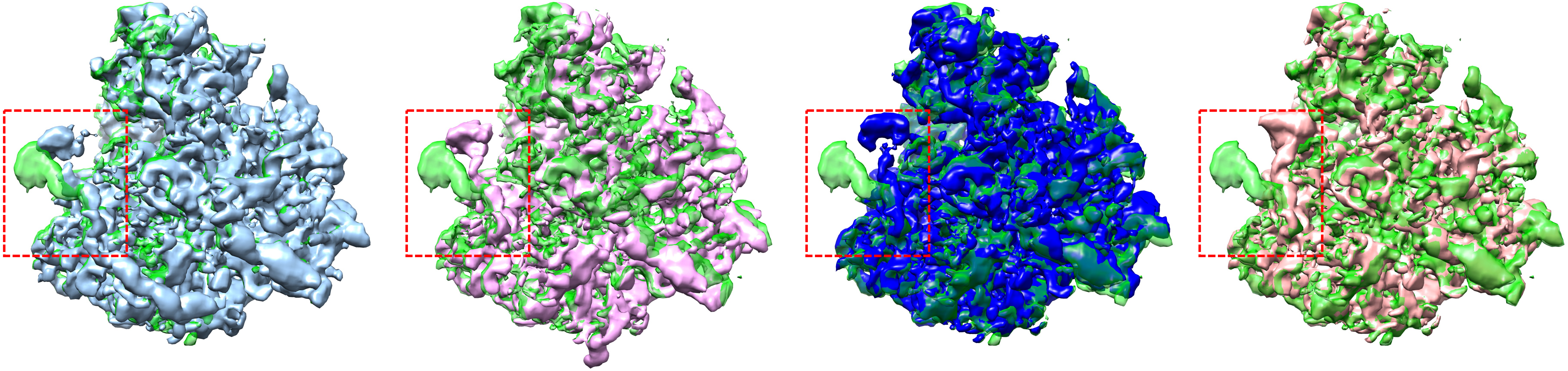

**Class 9 vs Class 8** **Class 9 vs Class 4** **Class 9 vs Class 6** **Class 9 vs Class 5**

**Fig.S4**

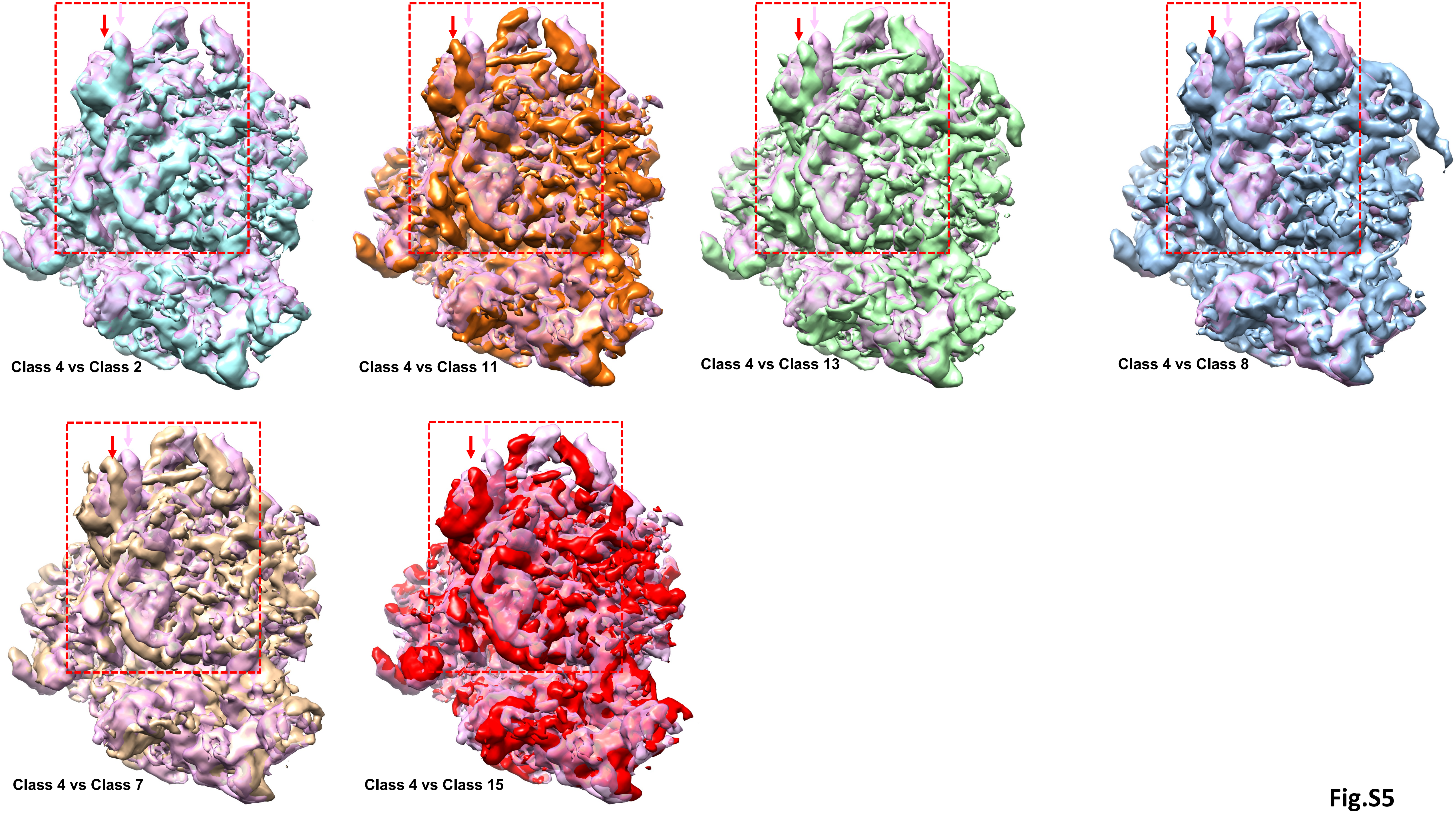
Class 4 vs Class 2
Class 4 vs Class 11
Class 4 vs Class 13
Class 4 vs Class 8
Class 4 vs Class 7
Class 4 vs Class 15

**Fig.S5**
